# Bridging the TB data gap: *in silico* extraction of rifampicin-resistant tuberculosis diagnostic test results from whole genome sequence data

**DOI:** 10.1101/628099

**Authors:** Kamela Charmaine S. Ng, Jean Claude S. Ngabonziza, Pauline Lempens, Bouke Catherine de Jong, Frank van Leth, Conor Joseph Meehan

**Author notes:** Corresponding Author: Kamela C S Ng, Nationalestraat 155, 2000 Antwerp, Belgium.

## Abstract

**Background:** *Mycobacterium tuberculosis* rapid diagnostic tests (RDTs) are widely employed in routine laboratories and national surveys for detection of rifampicin-resistant (RR)-TB. However, as next generation sequencing technologies have become more commonplace in research and surveillance programs, RDTs are being increasingly complemented by whole genome sequencing (WGS). While comparison between RDTs is difficult, all RDT results can be derived from WGS data. This can facilitate continuous analysis of RR-TB burden regardless of the data generation technology employed. By converting WGS to RDT results, we enable comparison of data with different formats and sources particularly for low and middle income high TB burden countries that employ different diagnostic algorithms for drug resistance surveys. This allows national TB control programs (NTPs) and epidemiologists to utilize all available data in the setting for improved RR-TB surveillance.

**Methods:** We developed the Python-based MTB Genome to Test (MTBGT) tool that transforms WGS-derived data into laboratory-validated results of the primary RDTs – Xpert MTB/RIF, XpertMTB/RIF Ultra, GenoType MDRTB*plus* v2.0, and GenoscholarNTM+MDRTB II. The tool was validated through RDT results of RR-TB strains with diverse resistance patterns and geographic origins and applied on routine-derived WGS data.

**Results:** The MTBGT tool correctly transformed the SNP data into the RDT results and generated tabulated frequencies of the RDT probes as well as rifampicin susceptible cases. The tool supplemented the RDT probe reactions output with the RR-conferring mutation based on identified SNPs. The MTBGT tool facilitated continuous analysis of RR-TB and Xpert probe reactions from different platforms and collection periods in Rwanda.

**Conclusion:** Overall, the MTBGT tool allows low and middle income countries to make sense of the increasingly generated WGS in light of the readily available RDT results, and assess whether currently implemented RDTs adequately detect RR-TB in their setting. With its feature to transform WGS to RDT results and facilitate continuous RR-TB data analysis, the MTBGT tool may bridge the gap between and among data from periodic surveys, continuous surveillance, research, and routine tests, and may be integrated within the existing national connectivity platform for use by the NTP and epidemiologists to improve setting-specific RR-TB control. The MTBGT source code and accompanying documentation is available at https://github.com/KamelaNg/MTBGT.

## Introduction

Resistance to rifampicin (RIF), the most potent anti-tuberculosis (TB) drug, hampers TB control. Rifampicin-resistant (RR)-TB persists as an urgent public health crisis as only 29% of estimated RR-TB patients world-wide were detected and notified in 2017. Further, 18% of previously treated TB patients were found to have RR-TB (WHO 2018). The WHO endorsed the following rapid molecular RR-TB diagnostic test (RDTs) to address this concern: Xpert MTB/RIF (Xpert Classic) and the new version Xpert MTB/RIF Ultra (Ultra) [Cepheid, Sunnyvale, USA] which employ heminested real-time polymerase chain reaction and molecular beacon technology (Blakemore et al. 2010); and the line probe assays – GenoType MDRTB*plus* v2.0 (hereinafter called LPA-Hain) [Hain Lifescience GmbH, Nehren, Germany]; and GenoscholarNTM+MDRTB II (hereinafter called LPA-Nipro) [NIPRO Corporation, Osaka, Japan] which rely on multiplex amplification and reverse hybridization of target to both wild-type and mutant probes on a membrane strip (Dheda et al. 2017). These tests were designed to detect RR-conferring mutations within the rifampicin resistance determining region (RRDR) or hotspot of the *rpoB* gene (Andre et al. 2017; Blakemore et al. 2010; Ng et al. 2018a; Ng et al. 2018c).

The Xpert assay involves binding of five short overlapping fluorescent probes to wild-type regions of the RRDR. Each of the five probes corresponds to several mutations which inhibit probe binding, thereby disrupting the signal from the respective probe. Thus, when a mutation is detected by Xpert, at least one probe does not bind. Certain mutations which completely interfere with probe binding result in ‘drop-out’ or absent probe, whereas mutations that allow limited probe hybridization are represented by ‘delayed’ probes (Blakemore et al. 2010). Xpert probes are therefore often considered as a proxy for circulating rifampicin resistance-conferring mutations which may be common or infrequent in a particular country. Understanding their frequency can be crucial for rifampicin resistance surveillance. For example, mutation Ser450Leu repeatedly detected globally, (Coll et al. 2018; Walker et al. 2015), is captured by ‘absent’ Xpert probe E (Ng et al. 2018a).

The implementation of Xpert was a breakthrough in TB diagnosis as it revolutionized detection of RR-TB worldwide, allowing for prompt identification of patients who need to undergo adapted treatment. Xpert Classic is the most widely deployed RDT globally, implemented as the initial diagnostic tool for all presumptive pulmonary TB patients by 32 out of 48 high TB-burden countries (WHO 2018). The wide utilization of the tests in both low and high burden TB countries resulted in the production of large volumes of RDT data, although comparisons within and between countries can be difficult due to use of differing technologies.

The utility of next generation sequencing technologies has been widely studied for improved detection of drug-resistant TB in diverse laboratory settings worldwide (Gardy & Loman 2018). Next generation sequencing-based whole genome sequencing (WGS) of *Mycobacterium tuberculosis* (*Mtb*) has been shown to accurately detect RR-TB by calling relevant single nucleotide polymorphisms (SNPs) in the *rpoB* gene (Coll et al. 2018; Miotto et al. 2017). WGS is already being widely implemented in the United Kingdom, the Netherlands, and New York, aimed towards completely replacing phenotypic drug susceptibility testing in the clinic (CRyPTICConsortium et al. 2018; de Viedma 2019). WGS implemented in high burden TB countries was shown to accurately estimate the prevalence of DR-TB (Zignol et al. 2018). The conventional periodic TB drug resistance surveys have been gathering data representative of the *Mtb* population in poor resource settings, while high income countries typically apply continuous surveillance (WHO 2018; Zignol et al. 2018), to help improve the choice of standard TB treatment before full drug sensitivity profile is known (WHO 2018).

The increasing use of WGS in research and public health initiatives can lead to a disconnect from the RDT-based data being generated routinely in the clinic, and the majority of surveys, widening the *Mtb* data gap. We aim to bridge this *Mtb* data gap by transforming WGS data into each of the related RDT data outputs, to facilitate analysis of RR-TB prevalence and underlying mutations regardless of the switch between data generation technologies. This will allow end-users to compare ‘apples with apples’, and analyze previous historical strains with current isolates representative of the entire TB patient population in the country.

We present the Myc TB Genome to Test (MTBGT) tool, a Python 3-based set of scripts that rapidly transform WGS data type to RDT and mutation reports. The modules automatically generate frequencies of rifampicin-susceptible (RS) and RR-TB samples detected and supplement the RDT probes output with the detected RR-conferring mutation in the format of ‘wild-type amino acid-codon number-mutant amino acid’.

## Materials & Methods

We developed the MTBGT tool, a Python 3-executable that converts *Mtb* WGS data in the form of variant call format (VCF) or MTBseq (Kohl et al. 2018) tab files into the most likely output that would be observed from Xpert Classic, Xpert Ultra, LPA-Hain, and LPA-Nipro based on previously validated work (Ng et al. 2018a; Ng et al. 2018c). The MTBGT tool can be accessed through https://github.com/KamelaNg/MTBGT.

### The MTBGT tool

The MTBGT tool can be run on any python-enabled operating system with no additional prerequisites. The MTBGT workflow is shown in Figure 1. The input is a folder of files derived from a SNP calling pipeline, either in standard raw VCF format or tab format as output from MTBseq. The MTBGT tool assumes the standard H37Rv NC000962.3 genome was used for calling these SNPs. If not, the user may remap the genome positions to the specific RR-TB-related codons using a tab delimited mapping file. An example of this tab-separated file is bundled with the tool. By default, the module will run all the RDTs on the input files.

**Figure 1.**
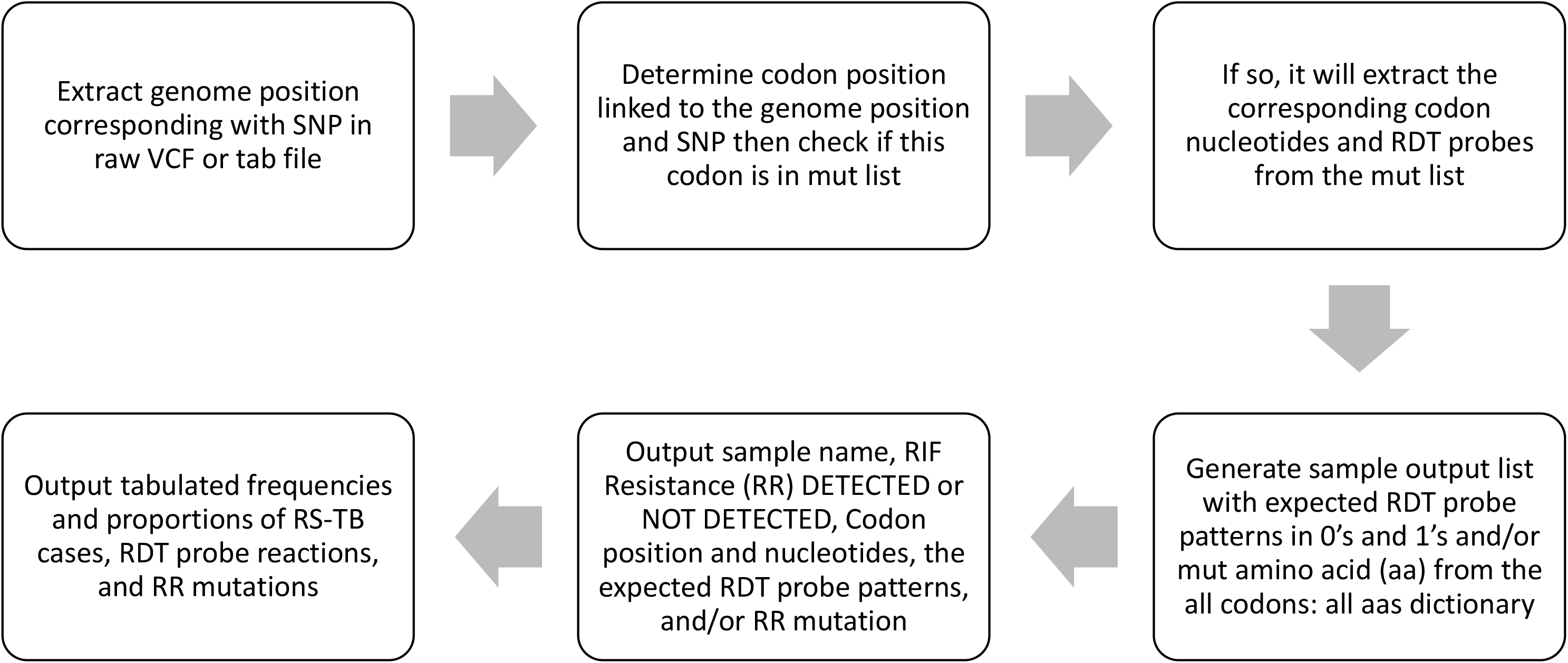
The MTBGT tool workflow.

The generated tab-delimited output file includes the Sample name, RIF resistance or susceptibility, the associated mutant codon position and mutation pattern, and a series of 0’s or 1’s indicating absence or presence of the capturing probe for the RDTs, and the RR-conferring mutation (An example output is given in Table 2). A summary table with counts and proportions of detected RS-TB cases, RDT probes, and RR-conferring mutations is also generated (Supplemental File Table S1). These tab-separated output files can be easily imported into Excel as outlined in the associated manual.

### Validation and sample application of the MTBGT modules

We simulated VCF files for internal validation of the MTBGT modules. We then randomly chose a VCF file and edited it in Notepad ++ 7.6.3 (https://notepad-plus-plus.org/) to contain all previously tested and validated RR-conferring mutations (Andre et al. 2017; Miotto et al. 2017) which were mapped to known RDT results (Ng et al. 2018a; Ng et al. 2018c) (Table 1). We generated files with single and multiple RR mutations to ensure the tool is robust for all scenarios. These simulated VCFs are provided with the tool.

**Table 1.**
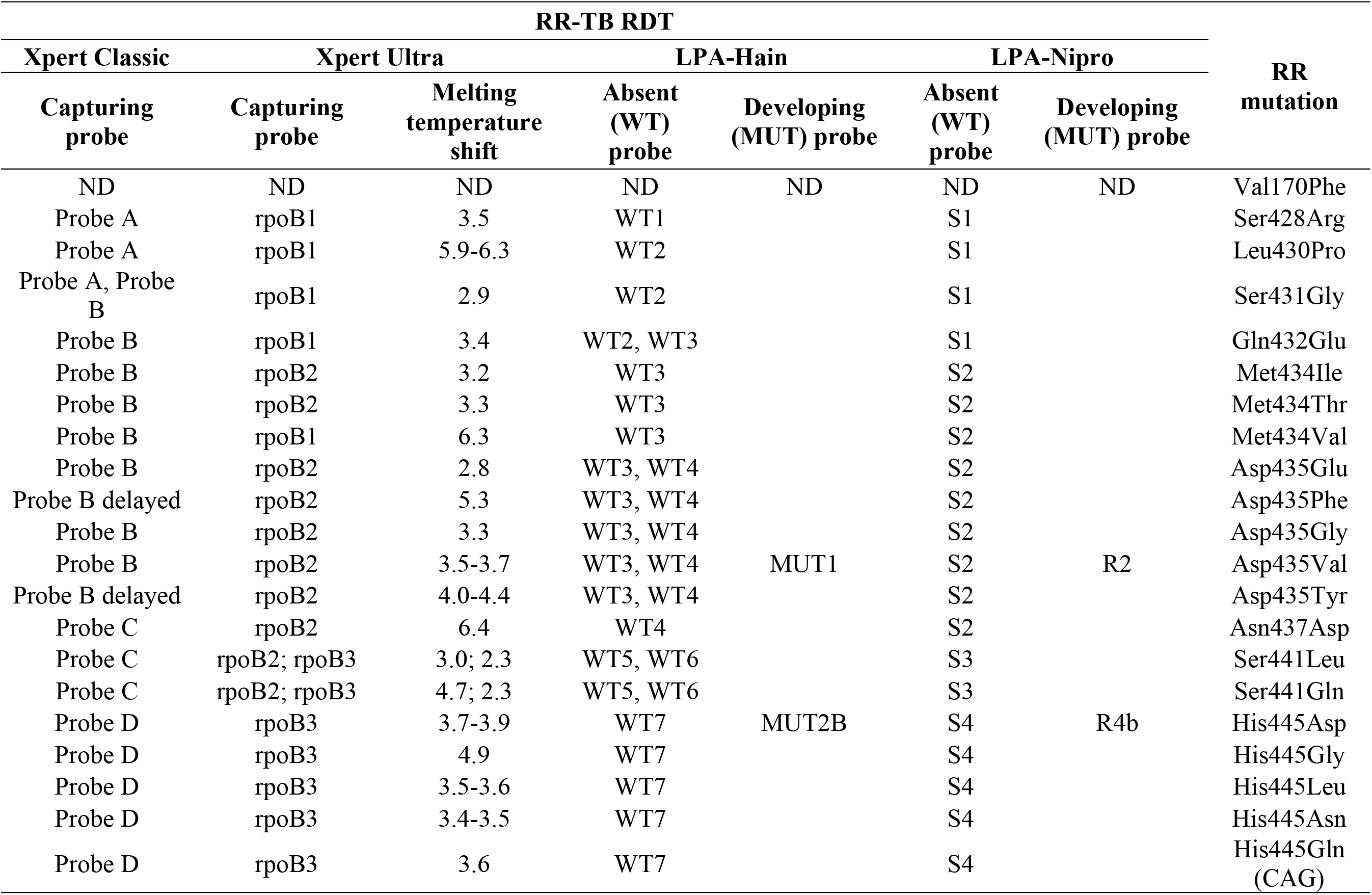
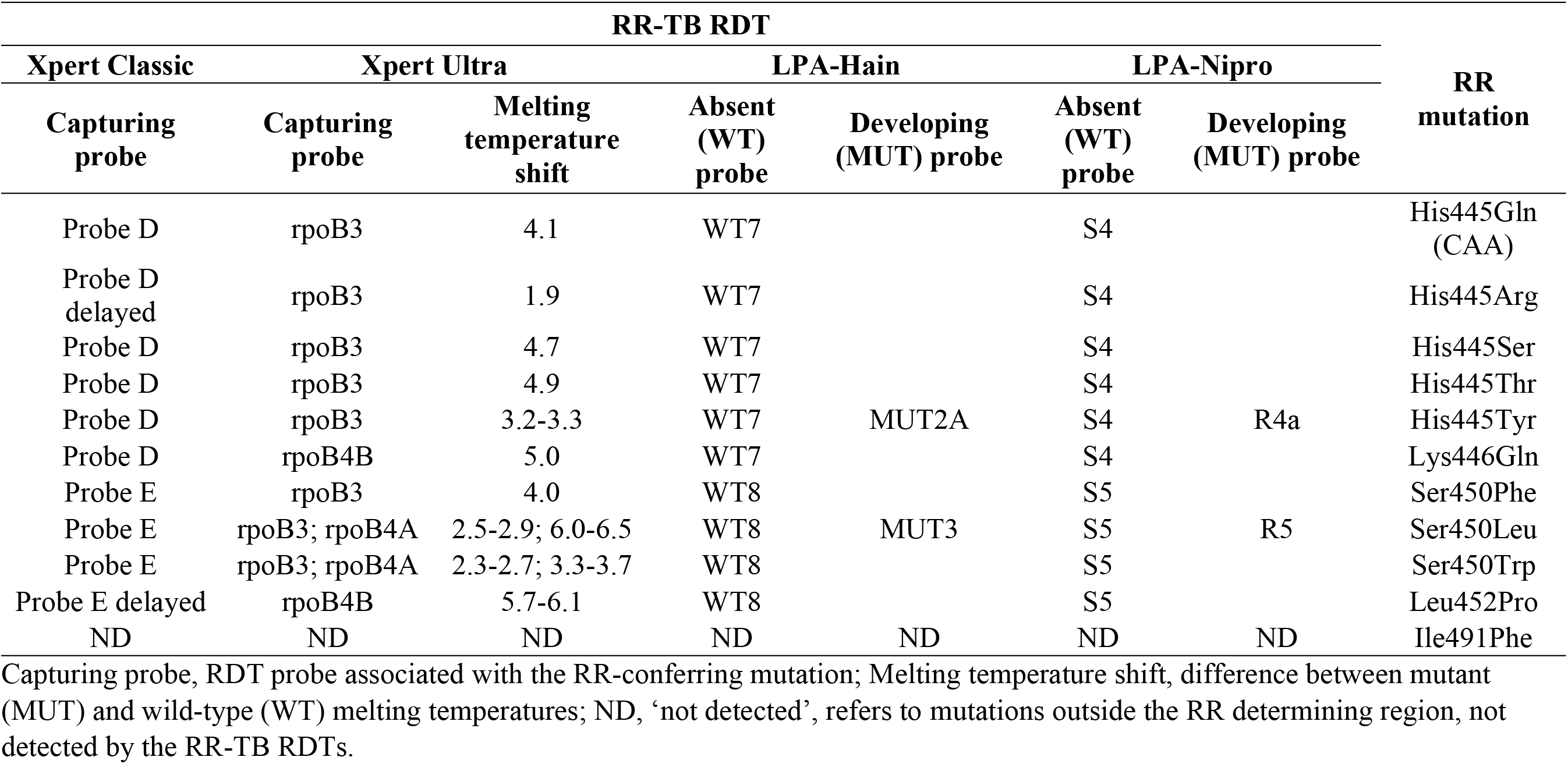
Rifampicin resistance (RR)-conferring mutations and associated rapid diagnostic test (RDT) results, previously tested and validated, and used as basis for developing the TBGT tool. Capturing probe, RDT probe associated with the RR-conferring mutation; Melting temperature shift, difference between mutant (MUT) and wild-type (WT) melting temperatures; ND, ‘not detected’, refers to mutations outside the RR determining region, not detected by the RR-TB RDTs.

To show real world applicability, the MTBGT tool was run on WGS of RS and RR-TB WHO Tropical Disease Research (TDR) strains with diverse resistance patterns and geographic origins stored in the Belgian Coordinated Collections of Microorganisms in the Institute of Tropical Medicine including the 47 TDR-TB strains tested in the previous validation of the RDTs against the available *rpoB* Sanger sequences of the strains (Ng et al. 2018a; Ng et al. 2018c; Vincent et al. 2012). The fastQ files (ENA accession PRJEB31023) for these samples were run through the MTBseq pipeline with default settings (Kohl et al. 2018) to generate the tab files for input to MTBGT. Additionally, WGS from 324 phenotypically RR-TB isolates of retreatment TB patients in Kinshasa, the Democratic Republic of Congo (DRC) collected in 2005-2010 (Meehan et al. 2018), and 233 WGS isolated 1991-2010 from Rwanda were subjected to the MTBseq pipeline and run through the MTBGT tool.

We were then able to extract the associated Xpert results from the 1991 to 2010 Rwandan dataset and compare it with actual Xpert results from 2012 to 2017 (Ng et al. 2018b).

## Results

We tested the MTBGT tool on a 64-bit Windows 10 Enterprise computer with a 2.50 GHz processor and a 8.00 GB of RAM. The running time was 39 milliseconds for 1 MTBseq tab file and 530 milliseconds for a VCF file.

The MTBGT tool correctly transformed the WGS-derived SNPs in the VCF and MTBseq tab files into the laboratory-validated RDT probe reactions (Table 2), and accurately detected all previously validated RR-conferring mutations in the simulated VCF and MTBseq clinical WGS data from the TDR, DRC (Figures 2A and 2B), and Rwanda (Figures 3A and 3B) strains, including double nucleotide changes – two SNPs covering two loci or genome positions – such as the CAC → AGC and CAC → TCC His445Ser mutations and combinations of any two unlinked mutations. The generated frequency and proportion tables (Supplemental File Table S1) showed the distribution of the RS and RR-TB strains, the RDT probe reactions, and the RR-conferring mutations detected.

**Table 2.** Example of combined results from the TBGT tool rapid diagnostic test (RDT) modules supplemented by the rifampicin resistance (RR)-conferring mutations detected.

| Filename | RIF Resistance | Codon number | Codon | Xpert Classic |  | Xpert Ultra |  | LPA-Hain |  | LPA-Nipro |  | RR mutation |
| --- | --- | --- | --- | --- | --- | --- | --- | --- | --- | --- | --- | --- |
|  |  |  |  | Capturing probe | Probe pattern | Capturing probe, Melting temperature shift | Probe pattern | Capturing probe | Probe pattern | Capturing probe | Probe pattern |  |
| DRC-052577 | DETECTED | 450 | TTG | Probe E | 1 1 1 1 0<br>1 | rpoB3,rpoB4 A; 2.5-2.9,6.0-6.5 | 1 1 0 0 1 | WT8, MUT3 | 1 1 1 1<br>1 1 1 0<br>0 0 0 1 | S5, R5 | 1 1 1 1<br>0 0 0 0<br>1 | Ser450Leu |
| DRC-091003 | DETECTED | 452 | CCG | Probe E delayed | 1 1 1 1 1<br>0 | rpoB4B; 5.7-6.1 | 1 1 1 1 0 | WT8 | 1 1 1 1<br>1 1 1 0<br>0 0 0 0 | S5 | 1 1 1 1<br>0 0 0 0<br>0 | Leu452Pro |
| 1993-09004 | DETECTED<br>(only by *rpoB* Sanger sequencing module) | 170 | TTC | NOT DETECTED |  | NOT DETECTED |  | NOT DETECTED |  | NOT DETECTED |  | Val170Phe |
| DRC-101308 | DETECTED<br>(only by *rpoB* Sanger sequencing module) | 491 | TTC | NOT DETECTED |  | NOT DETECTED |  | NOT DETECTED |  | NOT DETECTED |  | Ile491Phe |

**Figure 2.**
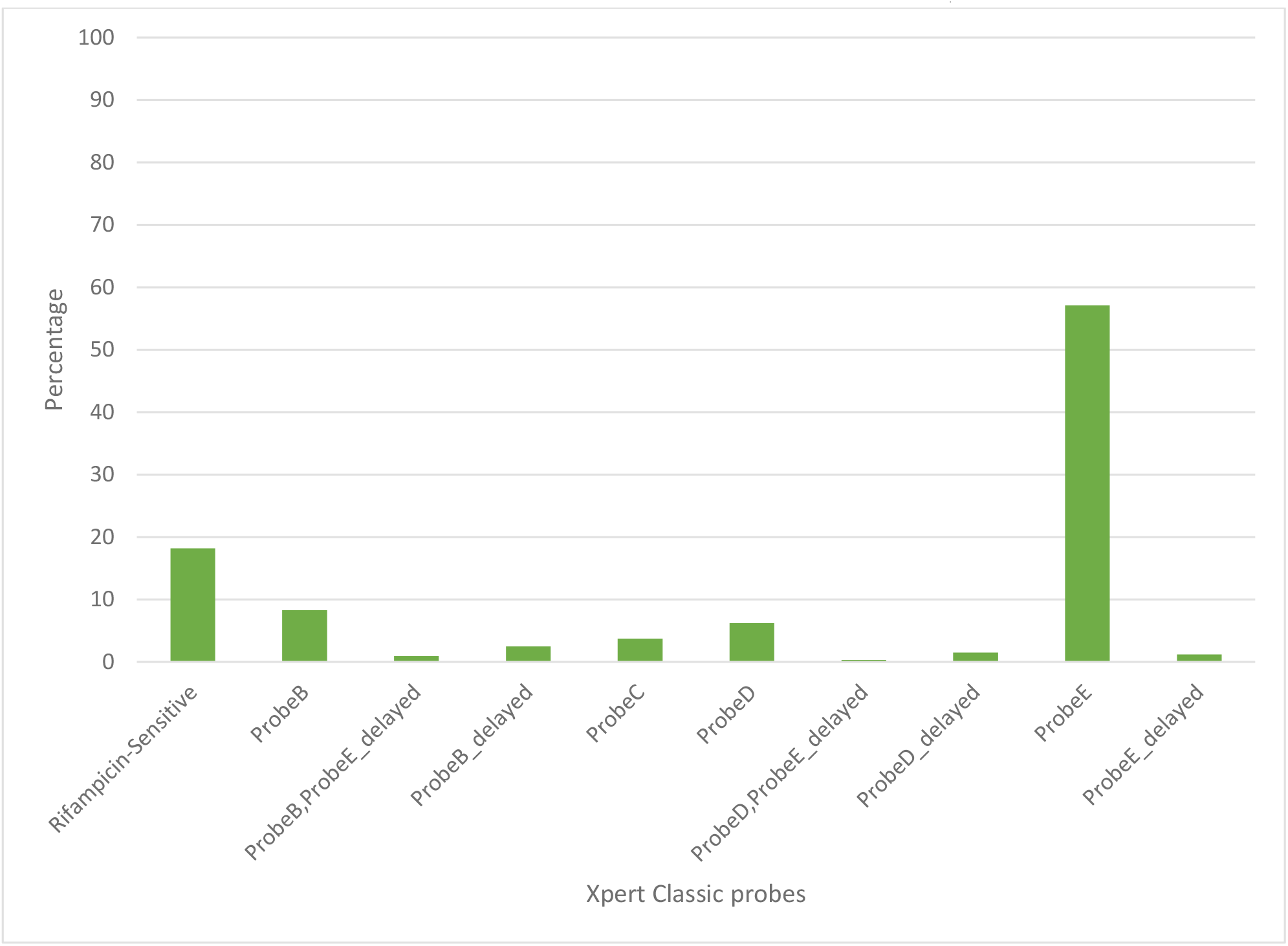
Distribution of rifampicin-sensitive samples and Xpert Classic probes among rifampicin-resistant tuberculosis isolates in Kinshasa, DRC from 2005 to 2010, detected by the MTBGT tool.

**Figure 3.**
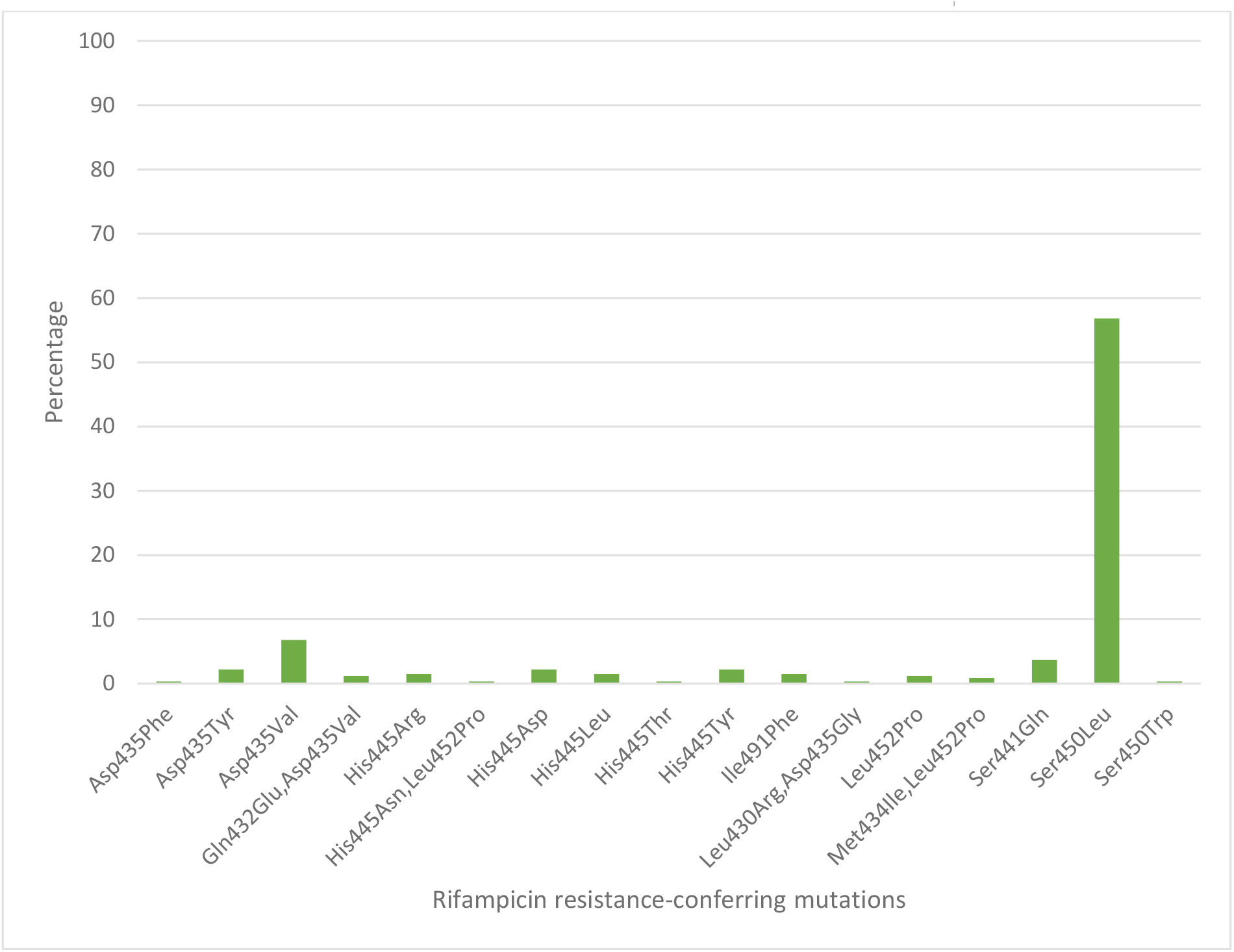
Distribution of rifampicin resistance-conferring mutations in Kinshasa, DRC from 2005 to 2010, detected by the MTBGT tool.

The *in silico* extracted Xpert results from the Rwandan WGS collected in 1991-2010 revealed that the majority of RR-TB cases detected were linked with Xpert Classic probe E (Figure 3A). This observation was consistent with the actual Xpert results gathered in 2012 to 2017. Figure 3C shows the distribution of 12 years worth of Xpert results in Rwanda. The MTBGT tool facilitated this continuous analysis of *in silico* predicted Xpert results from 2005 to 2010 WGS and actual Xpert results in 2012 to 2017, and allowed us to plot the distribution of absent probe E against the notified RR-TB cases from surveillance programs and national surveys in Rwanda. The low WGS sampling reflected in Figure 3C is explained by the limited TB cultures kept in the freezer in 2005 to 2010. Notably, we also observed the divergence of documented probe reactions from the predominant absent probe E in 2013 to 2017.

**Figure 4.**
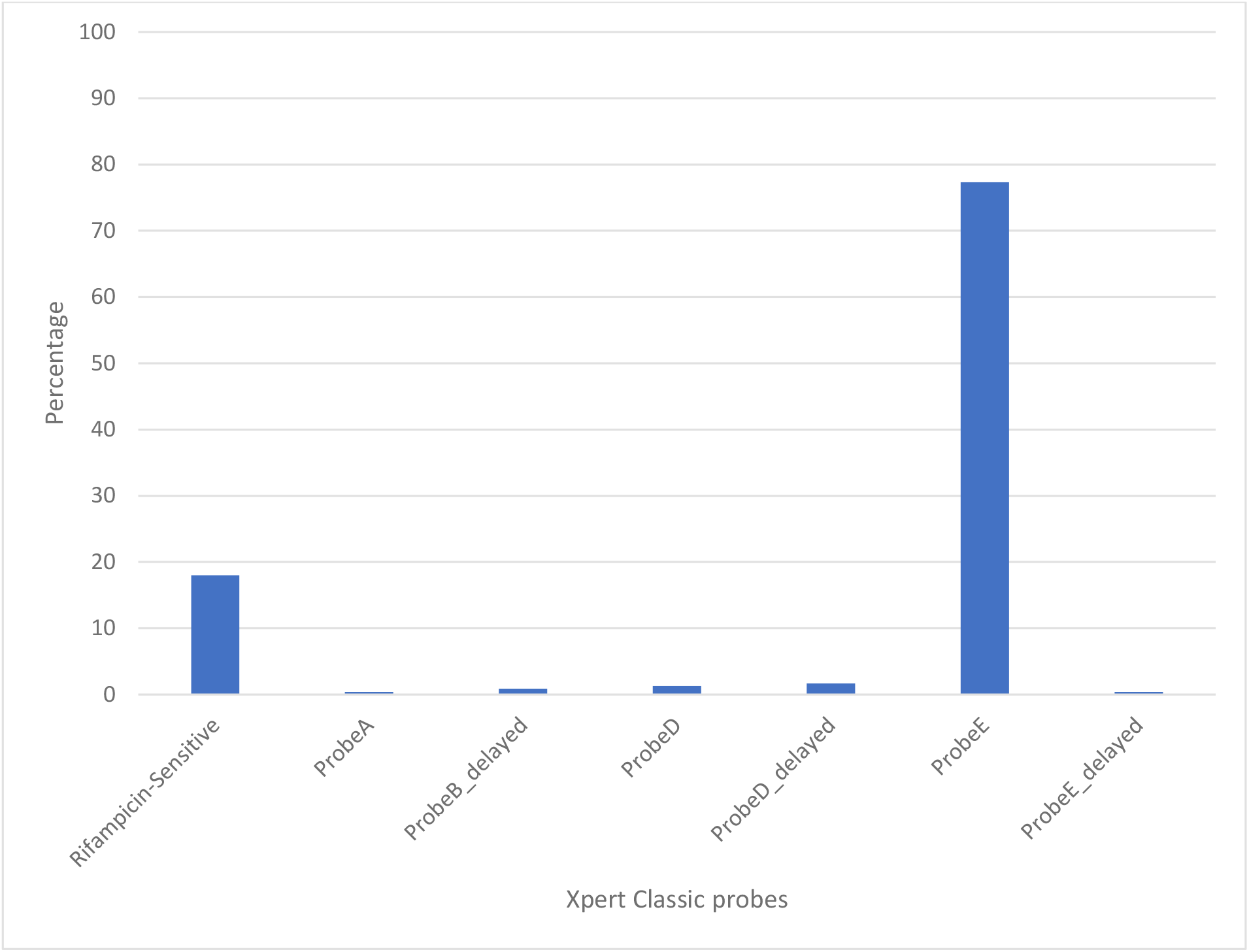
Distribution of rifampicin-sensitive samples and Xpert Classic probes among rifampicin-resistant tuberculosis isolates in Rwanda from 1991 to 2010, determined by the MTBGT tool.

Xpert Classic probe E was also predominantly seen in samples from Kinshasa, DRC (Figure 2A, Supplemental File Table S1). Mutation S450L, linked with Xpert Classic probe E, was the major RR-conferring mutation observed in both settings (Figures 2B and 3B).

**Figure 5.**
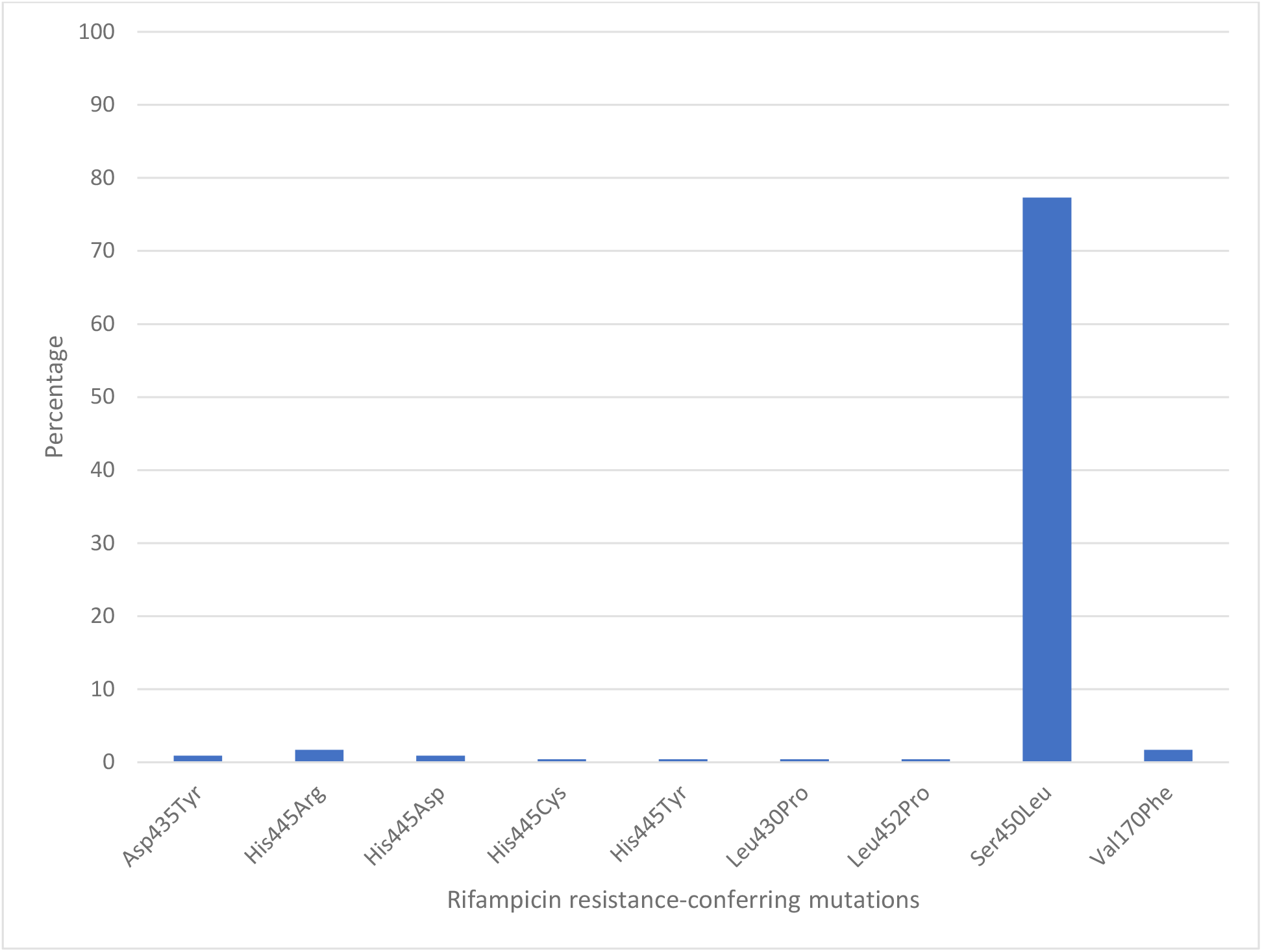
Distribution of rifampicin resistance-conferring mutations in Rwanda from 1991 to 2010, detected by the MTBGT tool.

## Discussion

To make sense of the increasingly produced WGS in light of the existing and readily available routine RDT results, we developed, validated, and applied the Myc TB Genome to Test (MTBGT), a Python-implemented tool that extracts RR-TB RDT results and RR-conferring mutations from WGS-derived SNP data.

If a country performed prior drug resistance surveys with a different diagnostic algorithm, the MTBGT tool allows for comparing data of different formats and sources, facilitating analysis of previous and current RR-TB case counts, probe reactions, and mutations, such as periodic DRS results conducted as cross-sectional surveys, often with different technology from the previous one. Although WGS data gives a much higher resolution than the RDTs, low and middle income high RR-TB burden countries are not yet capable of implementing routine WGS for all presumptive TB patients due to limitations in funding and logistic challenges (Meehan et al. 2019). However, RDTs are used routinely in such countries, producing a wealth of RR-TB data, whereas WGS is used for regular country-wide drug resistance surveys by the WHO (Zignol et al. 2018), and implemented in low burden high income countries. But since RDT results cannot be upscaled to WGS results, for comparison of these two data sources, WGS must be downscaled to the RDT results. By downscaling, the MTBGT tool adds to a substantial history of RDT probe data which amplifies surveillance analyses that can inform NTPs of low and middle-icome settings. This is similar to how CRISPR-based strain typing of *M. tuberculosis* (termed spoligotyping) can be predicted from WGS reads through SpoTyping (Xia et al. 2016). Although spoligotyping is known to have a lower resolution than WGS for typing, it is still widely used in many low and middle income TB endemic countries due to its low cost (Montoya et al. 2013; Suzana et al. 2017; Tulu & Ameni 2018). We foresee our tool being used in a similar manner, until WGS capabilities become more commonplace in low and middle income high RR-TB burden countries.

We show in Figure 3C how the MTBGT tool would be beneficial in Rwanda, a low income country which uses Xpert for routine diagnostics and WGS for WHO-led 5-year drug resistance surveys and selected research projects. Figure 3C reflects continuous distribution of RR-TB cases and Xpert absent probe E in Rwanda for more than a decade. These trends of RR-TB and absent probe E boost surveillance analyses that can inform the Rwandan National TB Control Program and support public health efforts for improved RR-TB control. The MTBGT tool allows for such comparisons and continuous distributions to be made, enabling low and middle income researchers to utilize all of their data for surveillance of RR-TB. This analysis of Rwandan datasets is a proof-of-concept that the MTBGT tool provides an opportunity to compare and analyze old and new data produced by different technologies bridging the RR-TB data gap. Care must be taken that the comparison is valid given the populations sampled. The predominant Xpert Classic probe E observed in Rwanda is supported by the associated RR mutation Ser450Leu (Tables 1 and 2) being present in the primary circulating multidrug-resistant TB clone in this setting (Ngabonziza et al. 2018), and also most frequently associated with global RR-TB (Cohen et al. 2015; Coll et al. 2018; Georghiou et al. 2016).

**Figure 6.**
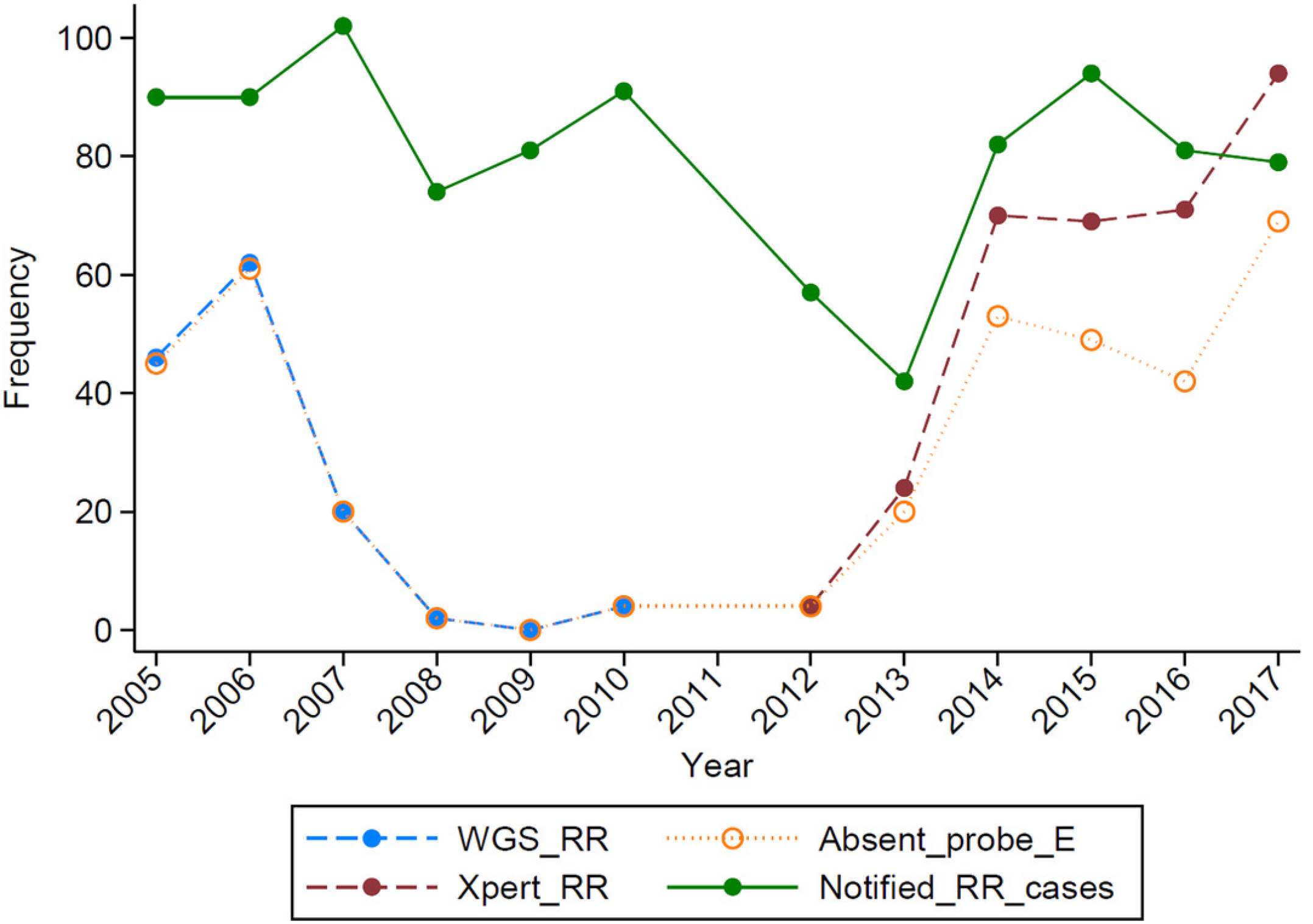
Distribution of RR-TB cases and Xpert Classic probe reactions predicted *in silico* from whole genomes collected in 2005 to 2010 in Rwanda and actual Xpert Classic results gathered in 2012 to 2017. The distribution of Xpert Classic absent probe E was overlaid on the documented RR-TB case counts and plotted against the notified RR-TB case counts from surveillance programs and national surveys in Rwanda.

Programatic surveillance is possible with only the rifampicin susceptibility status of patients’ samples, but at a different level. The RDT probe information which represents the underlying RR-conferring mutation provides an added critical value for epidemiological studies and surveillance (Ng et al. 2018b). This is exemplified by Xpert probe B and probe binding delay in South Africa being associated with false RR-TB results (Berhanu et al. 2019). In this setting, patients who were true rifampicin-susceptible (RS) but diagnosed as RR were treated with less effective and more toxic MDR drugs relative to the standard TB treatment. Further, non-routine WGS alone may give incorrect distribution of circulating mutations in the setting due to culture and sampling biases, as shown by the limited stored cultures resulting in low WGS sampling in Rwanda in 2005 to 2010.

Settings which employ different RDTs can refer to Table 1 where they can match laboratory-validated Xpert Classic and Xpert Ultra signatures per RR-TB mutation type (Ng et al. 2018a; Ng et al. 2018c). This would be helpful for settings such as South Africa which implements Xpert Classic and concurrently transitions to Xpert Ultra (Berhanu et al. 2018), and many other countries that will shift to Xpert Ultra in the near future, such as Rwanda by 2020.

The genome-based approach of the MTBGT tool also allows for reporting of disputed mutations that confer occult RR and are frequently missed by the *Mycobacterium* Growth Indicator Tube phenotypic DST (Ng et al. 2018a; Van Deun et al. 2015). For instance, disputed mutation Leu452Pro, epidemiologically linked with an extensively drug-resistant TB outbreak in KwaZulu-Natal, South Africa in 2005 (Cohen et al. 2015; Ioerger et al. 2009), was detected in some strains from Kinshasa, DRC and Rwanda (Figures 2B and 3B, Supplemental File). Mutation Leu452Pro is captured by delayed Xpert Classic probe E, denoting partially inhibited probe E fluorescence (Lawn & Nicol 2011; Ng et al. 2018a), and specifically identified by the unique combination of Xpert Ultra probe rpoB4B and corresponding melting temperature shift (Table 1), provided sufficient *Mtb* DNA is detected. Leu452Pro was reported to be missed in clinical samples by LPA-Hain due its end-probe location (Al-Mutairi et al. 2011; Rigouts et al. 2013), thus contributing to the RR-TB detection gap. The ability of the MTBGT tool to rapidly and accurately detect disputed mutations is therefore important.

As a supplementary feature, the MTBGT tool picks up any RR-conferring mutation present in the VCF or MTBseq tab file, and may help assess whether RDTs sufficiently detect RR-TB cases in specific settings.

Looking into the future when whole genome sequencing will be implemented as the primary diagnostic tool for all presumptive TB patients to confirm TB and/or RR-TB, the MTBGT tool will bridge the TB data gap by allowing comparison of past Xpert and LPA results with current *in silico* predicted results from routine whole genome sequences. These data would be comparable based on a common statistical parameter and representative of the presumptive TB population in the setting. With more whole genomes of *M. tuberculosis* sequenced, the *in silico* predicted probe distribution will more accurately capture the circulation of RR-TB mutations in the setting. The tool can do so by precisely revealing the divergence of current probes from a previously documented predominant probe over time, implicative of other circulating mutations such as that shown by actual Xpert results in 2013 to 2017 (Figure 3C).

The MTBGT tool was developed to aid surveillance efforts of low and middle income high RR-TB burden countries with likely fragmented data collection systems (Mazumdar et al. 2019). Through facilitated continuous analysis of previous historical and current RR-TB data, as well as clinical data from large aggregated files and routine laboratory and periodic survey RDT results combined with WGS-derived data from national surveillance programs and public data repositories, the MTBGT tool may create a larger picture of the RS-TB and RR-TB burden in a country. The RS and RR-TB counts and proportions and RDT probe reactions generated by the MTBGT tool may contribute to an extensive global database with years’ worth of data for continuous statistical modeling analyses and surveillance investigations.

Potentially, the MTBGT tool may be integrated in the national TB diagnostic algorithm through the existing connectivity platform or the newly implemented WHO cloud-based software (Dean 2019). The prospective application of the MTBGT tool may bridge and transform the *Mtb* data gap into action points for RR-TB clinicians to provide appropriate care for the individual TB/RR-TB patient, and the NTP, public health officials, and policy makers to intervene at the population-level for improved and sustained RR-TB control (Gardy & Loman 2018). The MTBGT modules could also be expanded to report non-RRDR RR-causing mutations and include other TB drugs – e.g. isoniazid, pyrazinamide, and fluoroquinolones.

## Conclusions

The MTBGT tool leverages improved access to next generation sequencing technologies in this genomic epidemiology era of TB, and complement *Mtb* WGS by rapidly transforming WGS files that store genomic sequence variations to validated outputs of the RR-TB RDTs. The prospective application of the MTBGT tool within a nationwide genomic epidemiology program will bridge the *Mtb* data gap among routine RDT data, research setting WGS, and periodic survey and continuous surveillance *rpoB* sequencing and WGS data. This may result in facilitated continuous analysis of the circulating RDT probes and underlying distribution of RR-conferring mutations in the setting, and may help assess whether currently implemented RDT(s) serve the detection of RR-TB cases in the country.

## Acknowledgements

We thank Button Ricarte for his insightful feedback on the manuscript.

## Additional Information & Declarations

### Data Availability

All scripts and data for validation may be accessed through GitHub: https://github.com/KamelaNg/MTBGT.

## References

Al-Mutairi NM, Ahmad S, and Mokaddas E. 2011. Performance comparison of four methods for detecting multidrug-resistant Mycobacterium tuberculosis strains. Int J Tuberc Lung Dis 15:110–115.

Andre E, Goeminne L, Cabibbe A, Beckert P, Kabamba Mukadi B, Mathys V, Gagneux S, Niemann S, Van Ingen J, and Cambau E. 2017. Consensus numbering system for the rifampicin resistance-associated rpoB gene mutations in pathogenic mycobacteria. Clin Microbiol Infect 23:167–172. 10.1016/j.cmi.2016.09.006

Berhanu RH, David A, da Silva P, Shearer K, Sanne I, Stevens W, and Scott L. 2018. Performance of Xpert MTB/RIF, Xpert Ultra, and Abbott RealTime MTB for Diagnosis of Pulmonary Tuberculosis in a High-HIV-Burden Setting. J Clin Microbiol 56. 10.1128/JCM.00560-18

Berhanu RH, Schnippel K, Kularatne R, Firnhaber C, Jacobson KR, Horsburgh CR, and Lippincott CK. 2019. Discordant rifampicin susceptibility results are associated with Xpert((R)) MTB/RIF probe B and probe binding delay. Int J Tuberc Lung Dis 23:358–362. 10.5588/ijtld.16.0837

Blakemore R, Story E, Helb D, Kop J, Banada P, Owens MR, Chakravorty S, Jones M, and Alland D. 2010. Evaluation of the analytical performance of the Xpert MTB/RIF assay. J Clin Microbiol 48:2495–2501. 10.1128/JCM.00128-10

Cohen KA, Abeel T, Manson McGuire A, Desjardins CA, Munsamy V, Shea TP, Walker BJ, Bantubani N, Almeida DV, Alvarado L, Chapman SB, Mvelase NR, Duffy EY, Fitzgerald MG, Govender P, Gujja S, Hamilton S, Howarth C, Larimer JD, Maharaj K, Pearson MD, Priest ME, Zeng Q, Padayatchi N, Grosset J, Young SK, Wortman J, Mlisana KP, O’Donnell MR, Birren BW, Bishai WR, Pym AS, and Earl AM. 2015. Evolution of Extensively Drug-Resistant Tuberculosis over Four Decades: Whole Genome Sequencing and Dating Analysis of Mycobacterium tuberculosis Isolates from KwaZulu-Natal. PLoS Med 12:e1001880. 10.1371/journal.pmed.1001880

Coll F, Phelan J, Hill-Cawthorne GA, Nair MB, Mallard K, Ali S, Abdallah AM, Alghamdi S, Alsomali M, Ahmed AO, Portelli S, Oppong Y, Alves A, Bessa TB, Campino S, Caws M, Chatterjee A, Crampin AC, Dheda K, Furnham N, Glynn JR, Grandjean L, Minh Ha D, Hasan R, Hasan Z, Hibberd ML, Joloba M, Jones-Lopez EC, Matsumoto T, Miranda A, Moore DJ, Mocillo N, Panaiotov S, Parkhill J, Penha C, Perdigao J, Portugal I, Rchiad Z, Robledo J, Sheen P, Shesha NT, Sirgel FA, Sola C, Oliveira Sousa E, Streicher EM, Helden PV, Viveiros M, Warren RM, McNerney R, Pain A, and Clark TG. 2018. Genome-wide analysis of multi- and extensively drug-resistant Mycobacterium tuberculosis. Nat Genet 50:307–316. 10.1038/s41588-017-0029-0

CRyPTICConsortium, the GP, Allix-Beguec C, Arandjelovic I, Bi L, Beckert P, Bonnet M, Bradley P, Cabibbe AM, Cancino-Munoz I, Caulfield MJ, Chaiprasert A, Cirillo DM, Clifton DA, Comas I, Crook DW, De Filippo MR, de Neeling H, Diel R, Drobniewski FA, Faksri K, Farhat MR, Fleming J, Fowler P, Fowler TA, Gao Q, Gardy J, Gascoyne-Binzi D, Gibertoni-Cruz AL, Gil-Brusola A, Golubchik T, Gonzalo X, Grandjean L, He G, Guthrie JL, Hoosdally S, Hunt M, Iqbal Z, Ismail N, Johnston J, Khanzada FM, Khor CC, Kohl TA, Kong C, Lipworth S, Liu Q, Maphalala G, Martinez E, Mathys V, Merker M, Miotto P, Mistry N, Moore DAJ, Murray M, Niemann S, Omar SV, Ong RT, Peto TEA, Posey JE, Prammananan T, Pym A, Rodrigues C, Rodrigues M, Rodwell T, Rossolini GM, Sanchez Padilla E, Schito M, Shen X, Shendure J, Sintchenko V, Sloutsky A, Smith EG, Snyder M, Soetaert K, Starks AM, Supply P, Suriyapol P, Tahseen S, Tang P, Teo YY, Thuong TNT, Thwaites G, Tortoli E, van Soolingen D, Walker AS, Walker TM, Wilcox M, Wilson DJ, Wyllie D, Yang Y, Zhang H, Zhao Y, and Zhu B. 2018. Prediction of Susceptibility to First-Line Tuberculosis Drugs by DNA Sequencing. N Engl J Med 379:1403–1415. 10.1056/NEJMoa1800474

de Viedma DG. 2019. Pathways and strategies followed in the genomic epidemiology of Mycobacterium tuberculosis. Infect Genet Evol. 10.1016/j.meegid.2019.01.027

Dean A. 2019. Sequencing for the surveillance of drug-resistant TB. WHO Global TB Programme.

Dheda K, Gumbo T, Maartens G, Dooley KE, McNerney R, Murray M, Furin J, Nardell EA, London L, Lessem E, Theron G, van Helden P, Niemann S, Merker M, Dowdy D, Van Rie A, Siu GK, Pasipanodya JG, Rodrigues C, Clark TG, Sirgel FA, Esmail A, Lin HH, Atre SR, Schaaf HS, Chang KC, Lange C, Nahid P, Udwadia ZF, Horsburgh CR, Jr., Churchyard GJ, Menzies D, Hesseling AC, Nuermberger E, McIlleron H, Fennelly KP, Goemaere E, Jaramillo E, Low M, Jara CM, Padayatchi N, and Warren RM. 2017. The epidemiology, pathogenesis, transmission, diagnosis, and management of multidrug-resistant, extensively drug-resistant, and incurable tuberculosis. Lancet Respir Med. 10.1016/S2213-2600(17)30079-6

Gardy JL, and Loman NJ. 2018. Towards a genomics-informed, real-time, global pathogen surveillance system. Nat Rev Genet 19:9–20. 10.1038/nrg.2017.88

Georghiou SB, Seifert M, Catanzaro D, Garfein RS, Valafar F, Crudu V, Rodrigues C, Victor TC, Catanzaro A, and Rodwell TC. 2016. Frequency and Distribution of Tuberculosis Resistance-Associated Mutations between Mumbai, Moldova, and Eastern Cape. Antimicrob Agents Chemother 60:3994–4004. 10.1128/AAC.00222-16

Ioerger TR, Koo S, No EG, Chen X, Larsen MH, Jacobs WR, Jr., Pillay M, Sturm AW, and Sacchettini JC. 2009. Genome analysis of multi- and extensively-drug-resistant tuberculosis from KwaZulu-Natal, South Africa. PLoS One 4:e7778. 10.1371/journal.pone.0007778

Kohl TA, Utpatel C, Schleusener V, De Filippo MR, Beckert P, Cirillo DM, and Niemann S. 2018. MTBseq: a comprehensive pipeline for whole genome sequence analysis of Mycobacterium tuberculosis complex isolates. PeerJ 6:e5895. 10.7717/peerj.5895

Lawn SD, and Nicol MP. 2011. Xpert(R) MTB/RIF assay: development, evaluation and implementation of a new rapid molecular diagnostic for tuberculosis and rifampicin resistance. Future Microbiol 6:1067–1082. 10.2217/fmb.11.84

Mazumdar S, Satyanarayana S, and Pai M. 2019. Self-reported tuberculosis in India: evidence from NFHS-4. BMJ Glob Health 4:e001371. 10.1136/bmjgh-2018-001371

Meehan CJ, Goig GA, Kohl TA, Verboven L, Dippenaar A, Ezewudo M, Farhat MR, Guthrie JL, Laukens K, Miotto P, Ofori-Anyinam B, Dreyer V, Supply P, Suresh A, Utpatel C, van Soolingen D, Zhou Y, Ashton PM, Brites D, Cabibbe AM, de Jong BC, de Vos M, Menardo F, Gagneux S, Gao Q, Heupink TH, Liu Q, Loiseau C, Rigouts L, Rodwell TC, Tagliani E, Walker TM, Warren RM, Zhao Y, Zignol M, Schito M, Gardy J, Cirillo DM, Niemann S, Comas I, and Van Rie A. 2019. Whole genome sequencing of Mycobacterium tuberculosis: current standards and open issues. Nat Rev Microbiol. 10.1038/s41579-019-0214-5

Miotto P, Tessema B, Tagliani E, Chindelevitch L, Starks AM, Emerson C, Hanna D, Kim PS, Liwski R, Zignol M, Gilpin C, Niemann S, Denkinger CM, Fleming J, Warren RM, Crook D, Posey J, Gagneux S, Hoffner S, Rodrigues C, Comas I, Engelthaler DM, Murray M, Alland D, Rigouts L, Lange C, Dheda K, Hasan R, Ranganathan UDK, McNerney R, Ezewudo M, Cirillo DM, Schito M, Koser CU, and Rodwell TC. 2017. A standardised method for interpreting the association between mutations and phenotypic drug resistance in Mycobacterium tuberculosis. Eur Respir J 50. 10.1183/13993003.01354-2017

Montoya JC, Murase Y, Ang C, Solon J, and Ohkado A. 2013. A molecular epidemiologic analysis of Mycobacterium tuberculosis among Filipino patients in a suburban community in the Philippines. Kekkaku 88:543–552.

Ng KC, Meehan CJ, Torrea G, Goeminne L, Diels M, Rigouts L, de Jong BC, and Andre E. (2018a. Potential Application of Digitally Linked Tuberculosis Diagnostics for Real-Time Surveillance of Drug-Resistant Tuberculosis Transmission: Validation and Analysis of Test Results. JMIR Med Inform 6:e12. 10.2196/medinform.9309

Ng KCS, Ngabonziza JCS, Meehan CJ, Migambi P, de Jong BC, Cobelens F, and van Leth F. (2018b. Automated algorithm for early identification of rifampicin-resistant tuberculosis transmission hotspots in Rwanda. The International Journal of Tuberculosis and Lung Disease 22:S1–S652.

Ng KCS, van Deun A, Meehan CJ, Torrea G, Driesen M, Gabriels S, Rigouts L, Andre E, and de Jong BC. (2018c. Xpert Ultra Can Unambiguously Identify Specific Rifampin Resistance-Conferring Mutations. J Clin Microbiol 56. 10.1128/JCM.00686-18

Ngabonziza JCS, Rigouts L, Mazarati JB, Uwizeye C, Nyaruhirira AU, Torrea G, de Jong BC, and Meehan CJ. 2018. Transmission drives the increase of multidrug-resistant tuberculosis in Rwanda. The International Journal of Tuberculosis and Lung Disease 22:S1–S652.

Rigouts L, Gumusboga M, de Rijk WB, Nduwamahoro E, Uwizeye C, de Jong B, and Van Deun A. 2013. Rifampin resistance missed in automated liquid culture system for Mycobacterium tuberculosis isolates with specific rpoB mutations. J Clin Microbiol 51:2641–2645. 10.1128/JCM.02741-12

Suzana S, Shanmugam S, Uma Devi KR, Swarna Latha PN, and Michael JS. 2017. Spoligotyping of Mycobacterium tuberculosis isolates at a tertiary care hospital in India. Trop Med Int Health 22:703–707. 10.1111/tmi.12875

Tulu B, and Ameni G. 2018. Spoligotyping based genetic diversity of Mycobacterium tuberculosis in Ethiopia: a systematic review. BMC Infect Dis 18:140. 10.1186/s12879-018-3046-4

Van Deun A, Aung KJ, Hossain A, de Rijk P, Gumusboga M, Rigouts L, and de Jong BC. 2015. Disputed rpoB mutations can frequently cause important rifampicin resistance among new tuberculosis patients. Int J Tuberc Lung Dis 19:185–190. 10.5588/ijtld.14.0651

Vincent V, Rigouts L, Nduwamahoro E, Holmes B, Cunningham J, Guillerm M, Nathanson CM, Moussy F, De Jong B, Portaels F, and Ramsay A. 2012. The TDR Tuberculosis Strain Bank: a resource for basic science, tool development and diagnostic services. Int J Tuberc Lung Dis 16:24–31. 10.5588/ijtld.11.0223

Walker TM, Kohl TA, Omar SV, Hedge J, Del Ojo Elias C, Bradley P, Iqbal Z, Feuerriegel S, Niehaus KE, Wilson DJ, Clifton DA, Kapatai G, Ip CLC, Bowden R, Drobniewski FA, Allix-Beguec C, Gaudin C, Parkhill J, Diel R, Supply P, Crook DW, Smith EG, Walker AS, Ismail N, Niemann S, Peto TEA, and Modernizing Medical Microbiology Informatics G. 2015. Whole-genome sequencing for prediction of Mycobacterium tuberculosis drug susceptibility and resistance: a retrospective cohort study. Lancet Infect Dis 15:1193–1202. 10.1016/S1473-3099(15)00062-6

WHO. 2018. Global tuberculosis report.

Xia E, Teo YY, and Ong RT. 2016. SpoTyping: fast and accurate in silico Mycobacterium spoligotyping from sequence reads. Genome Med 8:19. 10.1186/s13073-016-0270-7

Zignol M, Cabibbe AM, Dean AS, Glaziou P, Alikhanova N, Ama C, Andres S, Barbova A, Borbe-Reyes A, Chin DP, Cirillo DM, Colvin C, Dadu A, Dreyer A, Driesen M, Gilpin C, Hasan R, Hasan Z, Hoffner S, Hussain A, Ismail N, Kamal SMM, Khanzada FM, Kimerling M, Kohl TA, Mansjo M, Miotto P, Mukadi YD, Mvusi L, Niemann S, Omar SV, Rigouts L, Schito M, Sela I, Seyfaddinova M, Skenders G, Skrahina A, Tahseen S, Wells WA, Zhurilo A, Weyer K, Floyd K, and Raviglione MC. 2018. Genetic sequencing for surveillance of drug resistance in tuberculosis in highly endemic countries: a multi-country population-based surveillance study. Lancet Infect Dis 18:675–683. 10.1016/S1473-3099(18)30073-2

