## Supplementary material for "Bridging the TB data gap: *in silico* extraction of rifampicin-resistant tuberculosis diagnostic test results from whole genome sequence data": Summary tables generated by the TBGT tool (from another version of this preprint: version 1, 2019-05-05)

**Supplemental File**

Table S1. Summary tables of counts and proportions of rifampicin-susceptible TB samples, rapid diagnostic test probes, and rifampicin-resistant TB mutations detected by the TBGT tool from the 2005 to 2010 isolates in Kinshasa, DRC.

| Xpert classic probe | counts |
|---|---|
| ProbeB | 27 |
| ProbeB,ProbeE_delayed | 3 |
| ProbeB_delayed | 8 |
| ProbeC | 12 |
| ProbeD | 20 |
| ProbeD,ProbeE_delayed | 1 |
| ProbeD_delayed | 5 |
| ProbeE | 185 |
| ProbeE_delayed | 4 |
| Rif Sensitive | 59 |

| Xpert classic probe | proportions |
|---|---|
| ProbeB | 0.083 |
| ProbeB,ProbeE_delayed | 0.009 |
| ProbeB_delayed | 0.025 |
| ProbeC | 0.037 |
| ProbeD | 0.062 |
| ProbeD,ProbeE_delayed | 0.003 |
| ProbeD_delayed | 0.015 |
| ProbeE | 0.571 |
| ProbeE_delayed | 0.012 |
| Rif Sensitive | 0.182 |

| Xpert Ultra probe | counts |
|---|---|
| Rif Sensitive | 59 |
| rpoB1,rpoB2 | 4 |
| rpoB2 | 31 |
| rpoB2,rpoB3 | 12 |
| rpoB2,rpoB4B | 3 |
| rpoB3 | 25 |
| rpoB3,rpoB4A | 185 |
| rpoB3,rpoB4B | 1 |
| rpoB4B | 4 |

| Xpert Ultra probe | proportions |
|---|---|
| Rif Sensitive | 0.182 |
| rpoB1,rpoB2 | 0.012 |
| rpoB2 | 0.096 |
| rpoB2,rpoB3 | 0.037 |

| | |
|---|---|
| rpoB2,rpoB4B | 0.009 |
| rpoB3 | 0.077 |
| rpoB3,rpoB4A | 0.571 |
| rpoB3,rpoB4B | 0.003 |
| rpoB4B | 0.012 |

| Hain LPA probe | counts |
|---|---|
| Rif Sensitive | 59 |
| WT2,WT3,WT4,MUT1 | 4 |
| WT3,WT4 | 9 |
| WT3,WT4,MUT1 | 22 |
| WT3,WT8 | 3 |
| WT5,WT6 | 12 |
| WT7 | 11 |
| WT7,MUT2A | 7 |
| WT7,MUT2B | 7 |
| WT7,WT8 | 1 |
| WT8 | 5 |
| WT8,MUT3 | 184 |

| Hain LPA probe | proportions |
|---|---|
| Rif Sensitive | 0.182 |
| WT2,WT3,WT4,MUT1 | 0.012 |
| WT3,WT4 | 0.028 |
| WT3,WT4,MUT1 | 0.068 |
| WT3,WT8 | 0.009 |
| WT5,WT6 | 0.037 |
| WT7 | 0.034 |
| WT7,MUT2A | 0.022 |
| WT7,MUT2B | 0.022 |
| WT7,WT8 | 0.003 |
| WT8 | 0.015 |
| WT8,MUT3 | 0.568 |

| Nipro LPA probe | counts |
|---|---|
| Rif Sensitive | 59 |
| S1,S2,R2 | 4 |
| S2 | 9 |
| S2,R2 | 22 |
| S2,S5 | 3 |
| S3 | 12 |
| S4 | 11 |
| S4,R4a | 7 |
| S4,R4b | 7 |
| S4,S5 | 1 |

| | |
|---|---|
| S5 | 5 |
| S5,R5 | 184 |

| Nipro LPA probe | proportions |
|---|---|
| Rif Sensitive | 0.182 |
| S1,S2,R2 | 0.012 |
| S2 | 0.028 |
| S2,R2 | 0.068 |
| S2,S5 | 0.009 |
| S3 | 0.037 |
| S4 | 0.034 |
| S4,R4a | 0.022 |
| S4,R4b | 0.022 |
| S4,S5 | 0.003 |
| S5 | 0.015 |
| S5,R5 | 0.568 |

| Sanger probe | counts |
|---|---|
| Asp435Phe | 1 |
| Asp435Tyr | 7 |
| Asp435Val | 22 |
| Gln432Glu,Asp435Val | 4 |
| His445Arg | 5 |
| His445Asn,Leu452Pro | 1 |
| His445Asp | 7 |
| His445Leu | 5 |
| His445Thr | 1 |
| His445Tyr | 7 |
| Ile491Phe | 5 |
| Leu430Arg,Asp435Gly | 1 |
| Leu452Pro | 4 |
| Met434Ile,Leu452Pro | 3 |
| Rif Sensitive | 54 |
| Ser441Gln | 12 |
| Ser450Leu | 184 |
| Ser450Trp | 1 |

| Sanger probe | proportions |
|---|---|
| Asp435Phe | 0.003 |
| Asp435Tyr | 0.022 |
| Asp435Val | 0.068 |
| Gln432Glu,Asp435Val | 0.012 |
| His445Arg | 0.015 |
| His445Asn,Leu452Pro | 0.003 |
| His445Asp | 0.022 |

| | |
|---|---|
| His445Leu | 0.015 |
| His445Thr | 0.003 |
| His445Tyr | 0.022 |
| Ile491Phe | 0.015 |
| Leu430Arg,Asp435Gly | 0.003 |
| Leu452Pro | 0.012 |
| Met434Ile,Leu452Pro | 0.009 |
| Rif Sensitive | 0.167 |
| Ser441Gln | 0.037 |
| Ser450Leu | 0.568 |
| Ser450Trp | 0.003 |
